# Genetic analysis of coronary artery disease using tree-based automated machine learning informed by biology-based feature selection

**DOI:** 10.1101/2021.03.23.436652

**Authors:** Elisabetta Manduchi, Trang T. Le, Weixuan Fu, Jason H. Moore

## Abstract

Machine Learning (ML) approaches are increasingly being used in biomedical applications. Important challenges of ML include choosing the right algorithm and tuning the parameters for optimal performance. Automated ML (AutoML) methods, such as Tree-based Pipeline Optimization Tool (TPOT), have been developed to take some of the guesswork out of ML thus making this technology available to users from more diverse backgrounds. The goals of this study were to assess applicability of TPOT to genomics and to identify combinations of single nucleotide polymorphisms (SNPs) associated with coronary artery disease (CAD), with a focus on genes with high likelihood of being good CAD drug targets. We leveraged public functional genomic resources to group SNPs into biologically meaningful sets to be selected by TPOT. We applied this strategy to data from the UK Biobank, detecting a strikingly recurrent signal stemming from a group of 28 SNPs. Importance analysis of these uncovered functional relevance of the top SNPs to genes whose association with CAD is supported in the literature and other resources. Furthermore, we employed game-theory based metrics to study SNP contributions to individual level TPOT predictions and discover distinct clusters of well-predicted CAD cases. The latter indicates a promising approach towards precision medicine.

## 1 Introduction

In recent years, Machine Learning (ML) has gained increased appreciation as an alternative or complementary methodology to statistical approaches in ‘omics’ data analyses [1], [2], [3]. Setting up an appropriate ML pipeline for a given analysis task involves many decisions including data pre-processing algorithm selection, feature selection, feature engineering, estimator algorithm selection, and decisions about the many hyperparameter settings. Thus, of particular appeal are Automated ML (AutoML) methods, which assist (potentially non-expert) users in the design and optimization of ML pipelines [4]. Our group has developed a genetic programming-(GP-)based AutoML named Tree-based Pipeline Optimization Tool (TPOT) [5], [6], which has been successfully used to analyze data from metabolomics [7], [8], transcriptomics [9], [10], and toxicogenomics [10].

In addition to these ‘omics’ applications, an initial application of TPOT to a real-world genetic data set with prostate cancer aggressiveness as the endpoint discovered several feature combinations that significantly contributed to the classification accuracy [5]. The data set used in the latter was 1-2 orders of magnitude smaller, in terms of number of observations (∼2300 subjects), than the typical size of current Genome Wide Association Studies (GWAS). Moreover, biological filters suggested by the endpoint of interest were used to reduce the number of features to the manageable size of ∼200 Single Nucleotide Polymorphisms (SNPs). Even with this biological guidance, the predictive performance was much lower than that achieved in the other TPOT ‘omics’ applications cited above. This is, however, not surprising considering the challenges associated with complex trait GWAS data, such as missing heritability, typically small effect sizes of common variants, and genetic heterogeneity (i.e. different SNPs being responsible for the trait in different subjects) [11], [12].

In this work, we set to further explore both the challenges and potential insights of TPOT analyses on a largescale genotype data set via a case study in Coronary Artery Disease (CAD) leveraging the UK Biobank resource [13]. We note that, despite numerous large-scale GWAS, less than half of the of CAD heritability has been accounted for [14]. After identifying cases and controls, to establish a baseline, we first assessed the predictive performance of models using GWAS main effect CAD SNPs as features, i.e. SNPs previously identified from traditional univariate GWAS analyses. We then explored SNPs mapped to six genes suggested for CAD drug repurpos-ing and drug development [15]. For each of these genes, we also looked at SNPs mapping to its connected genes from Hetionet (https://het.io/), an integrative network of biomedical knowledge. For each gene in this extended network, we considered not only the SNPs in the gene body and proximal promoter region but also those residing in its potential enhancers based on publicly available epigenomic data from CAD relevant tissues (Figure 1A). We note that TPOT inherently analyzes features (i.e. SNPs) as a group and makes no assumptions about additivity of their effects. By utilizing a 2-stage TPOT approach and leveraging biologically meaningful SNP groupings, we identified a strikingly recurrent signal stemming from models built on an input subset of 28 SNPs. After ranking these SNPs according to permutation feature importance, we uncovered links between the top SNPs and other genes related to atherosclerotic plaques and myocardial infarction. We also analyzed contributions to the individual predictions for the 28 SNPs using Shapley Additive exPlanations (SHAP) [16]. We clustered cases which were well predicted by the optimal model based on SHAP values, aiming at dissecting their heterogeneity in terms of driver SNPs. This also highlighted specific groups of cases for whom the predictions were driven by SNPs mapping to genes whose CAD relevance is supported by the literature and other functional genomic resources. These results provide new hypotheses about the genetic basis of CAD and demonstrate the utility of AutoML for genetic association analysis as well as the potential of applying metrics such as SHAP to ML models for precision medicine studies.

**Fig. 1.**
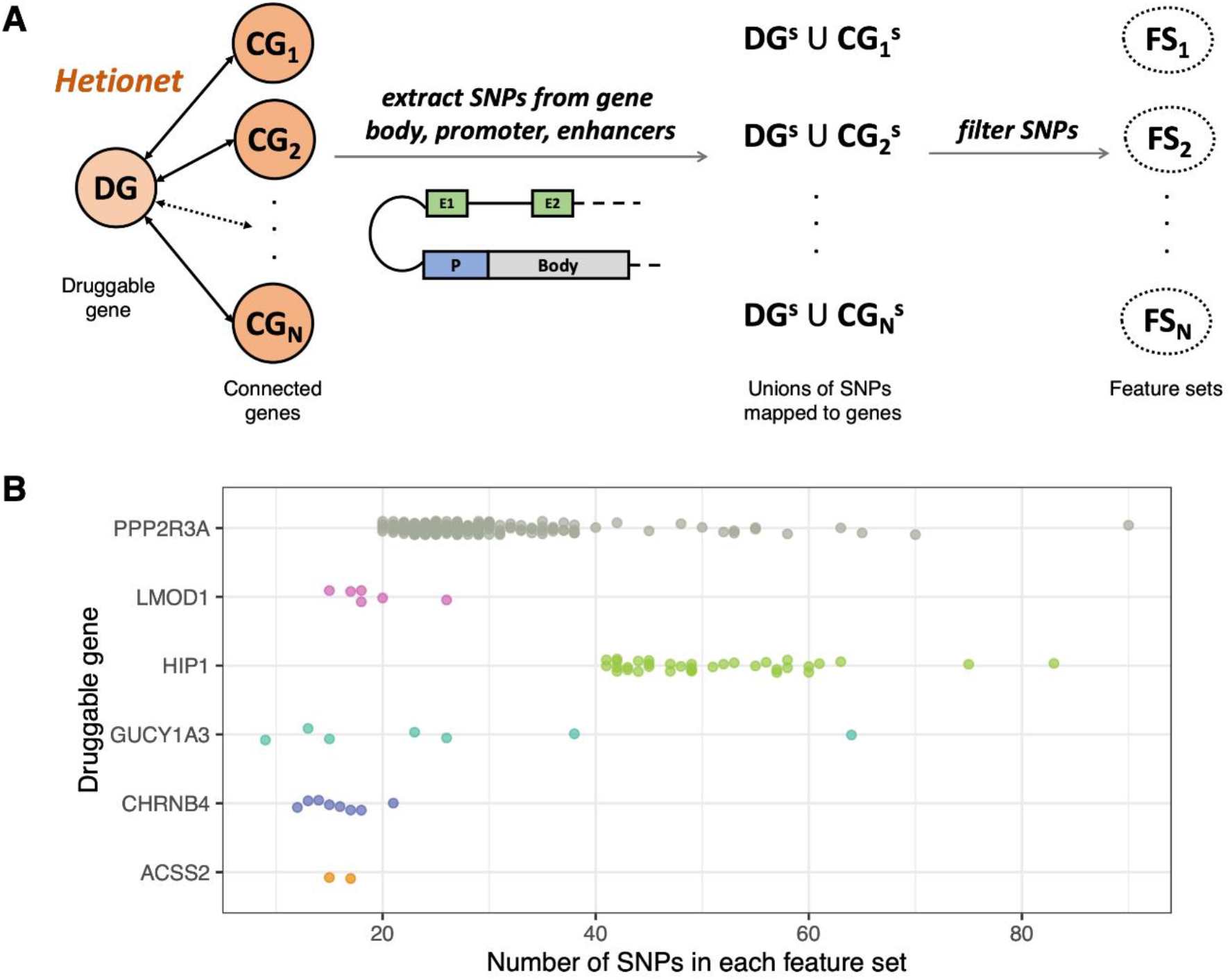
(A) Selection of SNPs and FSs. For each druggable gene DG, its connected genes (CGs) were obtained from Hetionet. For each CG, the SNPs mapping to its body, promoter, and putative enhancers were identified (CG^S^) and added to those mapping to DG (DG^S^). The corresponding FS was derived from these SNPs after filtering by functionality scorers and pruning. (B) FSs for the druggable genes. Each point corresponds to an FS for the DG indicated on the y-axis and its x-coordinate indicates the number of SNPs in that FS.

## 2 Methods

### 2.1 GWAS Data Preparation

From the UK Biobank (UKB) data, we extracted all subjects of white British ancestry (i.e. with a value of 1 for UKB field #22006) and retained a maximal subset of unrelated individuals (exploiting the related pairs file provided by UKB) whose genetically inferred sex matched the sex information collected at recruitment. We applied several filters based on flags in the following UKB fields: 22010 (recommended genomics exclusions), 22051 (UKBiLEVE quality control failure), 22019 (sex chromosome aneuploidy), 22021 (kinship inferences), 22027 (outliers for heterozygosity or missing rate). We defined CAD cases based on the criteria from Supplemental Table 1 of [17], arriving at a collection of 19,134 cases and 321,881 controls. For each such subject, we obtained the first 10 genetic Principal Components (PC) from UKB as well as the genotyping array and age. We defined age as the value at the last assessment center visit for individuals in the control group and at diagnosis/operation/death for individuals in the case group, depending on the field contributing to their case classification.

### 2.2 SNP Selection and Groupings

Our starting point were the six CAD ‘druggable’ genes suggested by [15] for drug repurposing (*CHRNB4, ACSS2*, and *GUCY1A3*) and drug development (*LMOD1, HIP1*, and *PPP2R3A*). We then obtained all autosomal genes connected with each druggable gene from Hetionet (https://het.io/), an integrative network assembling the knowledge from 29 different databases of genes, compounds, diseases, and more. For each gene (whether druggable or connected to a druggable gene), we obtained its GRCh37 coordinates from Ensembl genes 101 [18], extending them to include 5kb upstream and 1kb downstream of the Transcription Start Site. In addition, for each gene, we obtained its putative enhancers in CAD relevant tissues (fat, heart, and vascular) from the Roadmap Epigenomics Enhancer-Gene Links (https://ernstlab.biolchem.ucla.edu/roadmaplinking/). We then used BEDTools v2.25.0 [19] to extract, for each gene, SNPs residing in its body, promoter, or any of its enhancers (we only considered SNPs with a Minor Allele Frequency (MAF) > 0.01 and an imputation info score > 0.9). We further filtered the resulting collection of SNPs using software aimed at scoring their potential functionality (whether coding or non-coding). Namely, we used CADD [20] v1.6, GWAVA [21] v1.0, and TraP [22] v3.0, and we only retained SNPs satisfying at least one of these conditions: (1) CADD scaled score ≥ 10, or (2) GWAVA score ≥ 0.5, or (3) TraP score ≥ 0.459. Finally, for each druggable gene, we took the filtered SNPs mapping to the gene or any connected gene and pruned them for Linkage Disequilibrium (LD) using qctool v2 (https://www.well.ox.ac.uk/~gav/qctool_v2/) with a threshold of 0.8 for r2. For each druggable gene, we defined one Feature Set (FS) per connected gene, consisting of all SNPs resulting from the above filters and mapping either to the druggable or the connected gene (body, promoter, or any enhancer), as illustrated in Fig 1. Note that any two FSs of a druggable gene share all the SNPs of that gene.

### 2.3 TPOT Runs

In our first set of analyses, we used classic TPOT, whose source code is freely available at https://github.com/EpistasisLab/tpot. We then assessed the results derived by incorporating covariate adjustments as described in [10] using resAdj TPOT, whose code is also freely available at https://github.com/EpistasisLab/tpot/tree/v0.11.1-resAdj. In the latter analysis, we adjusted the outcome for age, sex and the first 10 PCs and we adjusted all features for genotyping array and the first 10 PCs. In all TPOT runs, we applied 5-fold cross validation. For each TPOT run, to match 19,134 cases, we randomly selected 19,134 samples from 321,881 individuals in the control group to obtain a balanced and reasonably sized input dataset. Where specified (see Results section), we used the Template and Feature Set Selector (FSS) described in [9]. The Template constrains the GP to only examine pipelines with a given architecture. The FSS slices the input data set into smaller sets of features, allowing the GP to select the best subset in the final pipeline.

### 2.4 Feature Assessments

We employed ELI5 v0.10.1 (https://github.com/TeamHG-Memex/eli5) to calculate permutation feature importance and the python SHAP library (https://github.com/slundberg/shap) to compute SHAP values with kernel explainer, an agnostic method that makes no assumption on the model type. Moreover, we used shap.kmeans to generate the explainer background from the training set, with 46 clusters for classic TPOT and 73 for resAdj TPOT. We arrived at these numbers by examining the Dunn indices for *k*-means clusterings for *k* varying between 30 and 100 and selecting the *k* yielding the highest value, using the R package NBClust [23]. The Dunn index is a measure of cluster quality defined in [24]. We also employed NbClust to inspect Dunn indices and generate *k*-means clustering of subjects based on SHAP values.

For visualization, we used the python SHAP library to compute a matrix of SHAP values for each individual and SNP and produce the initial summary plots. From the SHAP value matrix, we generated the final force plots using the R programming language (v 4.0.3) with the dplyr (v1.0.2), ggplot2 (v3.3.2), tidyr (v1.1.2), readr (v1.4.0), and seriation (v1.2-9) libraries. A GitHub repository with reproducible R visualization code is available at https://github.com/trang1618/cad-shap.

## 3 Results

We first explored the predictive performance of TPOT when using as features the SNPs in the CAD loci identified in [25] and reported in Supplemental Table 2 of that paper. After LD pruning (with a threshold of 0.6 for r2) we obtained 92 SNPs. We ran classic TPOT 50 times (without a Template), each with a random down-sampling of the controls (hence with a balanced input of 19,134 cases and 19,134 controls). In each run, the input was split into training (75%) and holdout validation testing (25%) parts. We set a population of 100 in the GP and the stopping criterion was the earliest of 100 generations or 2 days. Over the 50 runs, the range for the accuracy of the TPOT optimized pipeline on the holdout testing set was 0.561-0.582, which is reasonable given the typically small effect sizes of common variants and genetic heterogeneity. This classic TPOT result served to establish a reference based on the strongest known main effect signals, suggesting that runs that explore other sets of variants may not yield accuracy values much higher than 0.50. Therefore, particularly useful for this type of application, is TPOT’s feature set selector (FSS), which slices the input data set into smaller sets of features and reports the best feature subset in the final pipeline. Indeed, due to the GP stochasticity, we carried out multiple runs of TPOT. We could therefore examine the consistency with which the same FS was selected across multiple runs, hypothesizing that an FS repeatedly selected in different pipelines contains potentially interesting signals.

To look for variants other than the known GWAS hits, we focused on SNPs mapped to the body, promoter, or putative enhancers (in CAD relevant tissues) of the six ‘druggable’ genes from [15] and their connected genes from Hetionet, as described in Methods. Since runs of TPOT on such large data sets are computationally expensive, we first carried out pilot analyses consisting of 10 TPOT runs per druggable gene, using the Template *FSS-Transformer-Classifier*, where each FS comprised the SNPs mapped to a druggable gene and one of its connected genes (see Methods and Fig. 1). In these pilot analyses we had the same settings as the above baseline analyses in terms of down-sampling of controls, train/test split, and GP population, but the GP stopping criterion was shortened to the earliest of 100 generations or 1 day. The best out of 10 testing accuracies for the six druggable gene varied from 0.5062 (for the runs using FSs derived from *LMOD1*) to 0.5229 (for the runs using FSs derived from *PPP2R3A*). Moreover, out of 197 FSs considered for *PPP2R3A*, the same FS (corresponding to its connected gene *PRC1*) was selected in 4 of the 10 runs. Thus, we decided to pursue the 197 FSs determined by *PPP2R3A* and its connected genes for more extensive TPOT runs, as we had indication of possible interesting signals among these SNPs.

In the more extensive runs, we adopted a 2-stage procedure, illustrated in Fig. 2. In both stages we increased the number of runs from 10 to 50. Moreover, we made sure to use different holdout testing sets in the two stages. More precisely, we generated 50 random down-samplings of the controls and, for each of these, we randomly split the resulting 38,264 cases and controls into a training (75%) part, a holdout validation testing (13%) part for stage 1 and a holdout validation testing part (12%) for stage 2. In the first stage, we carried out 50 TPOT runs (one for each of the down-sampling and train/test splits), using the Template *FSS-Transformer-Classifier*, focusing only on the FSs corresponding to *PPP2R3A*. We used a population of 100 in the GP and a stopping criterion of the earliest of 100 generations or 1 day. We noted that in 21 of the 50 runs the TPOT optimized pipeline selected the same FS, corresponding to *PRC1*, re-enforcing the results from the pilot runs. Moreover, out of the 50 runs, the accuracy of the best pipeline on the holdout testing set was 0.5245 and this pipeline selected the *PRC1* feature set (consisting of 28 SNPs mapped to either this gene or *PPP2R3A*).

**Fig. 2.**
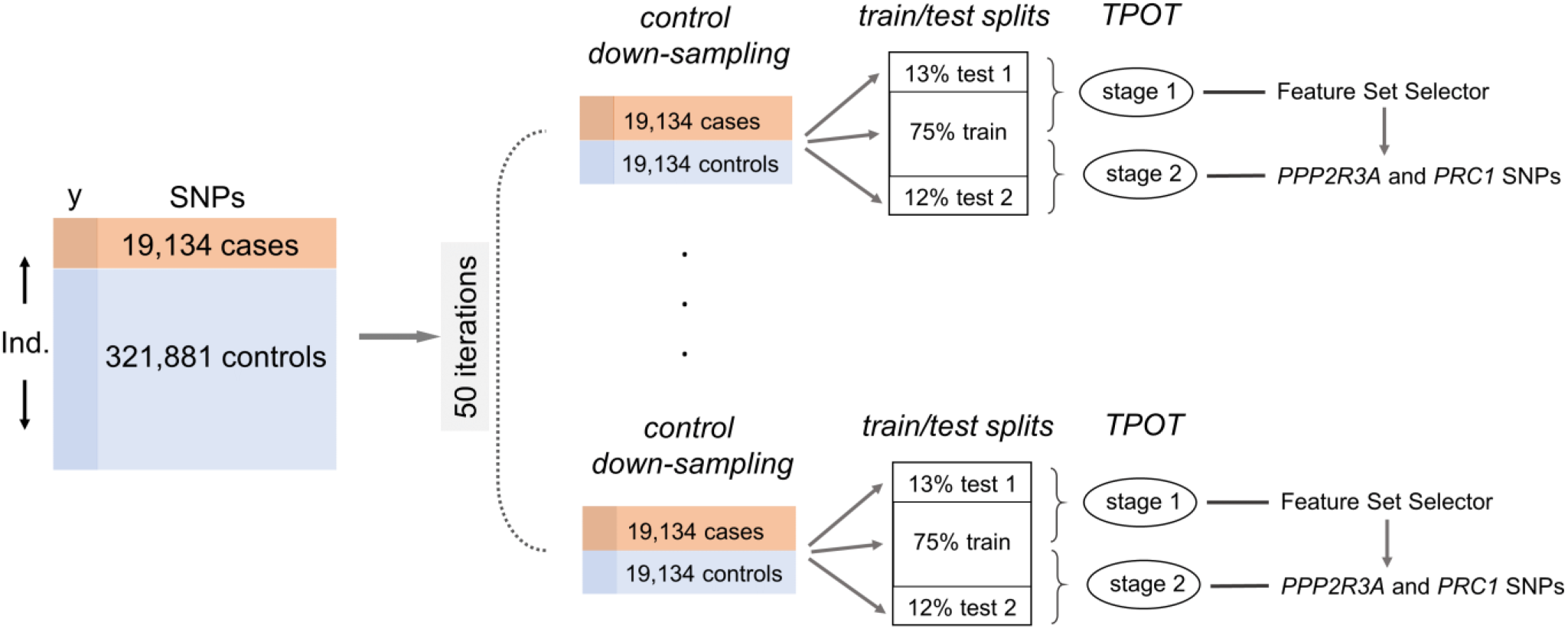
Workflow for the 2-stage procedure.

To assess the significance of these results, we performed permutation tests. Ideally, we would generate 1000 permutations and, for each permutation, repeat the entire stage 1 procedure of 50 runs set up as above. However, because of the computationally expensive nature of GP, we only generated 20 permutations of the target column in our full data set and repeated the stage 1 analysis in each permutation (for a total of 20×50=1000 TPOT runs). For each of the 20 permutations, we investigated the highest occurrence frequency of the same FS out of 50 runs and all of these 20 values were ≤ 4, much smaller than the observed value of 21 on the unpermuted data. We also assessed the highest testing accuracy out of 50 runs for each of these permutations, and all of these 20 values were less than the observed value (0.5245) on the unpermuted data. Even though we cannot infer that the results from stage 1 have permutation p-values < 0.05 due to the limited number of permutations, it is nevertheless noteworthy to see that the same FS was selected in such a large proportion of the 50 original TPOT runs compared to the best proportions achieved in the runs on the 20 permuted data sets.

In stage 2, we focused on the SNPs from the significant FS from stage 1, i.e. the 28 SNPs mapped to either *PPP2R3A* or *PRC1* (see Supplemental Table 1) and ran TPOT without Template and extending the stopping criterion to the earliest of 100 generations or 2 days, to see if we could improve accuracy. Again, we ran TPOT 50 times using the down-sampling and train/test splits illustrated above, but this time, the accuracies were computed using the holdout testing sets 2. These unconstrained runs slightly improved the testing accuracy, with the best of 50 accuracies equal to 0.5274. For each of the 20 permutations, we ran 50 TPOT runs with the stage 2 settings (again, for a total of 1000 runs) and again the highest testing accuracy out of 50 runs was less than the observed value across all permutations.

The classic TPOT results indicate that there is signal within the combination of SNPs mapped to the body/promoter/enhancers of *PPP2R3A* and *PRC1*. In order to verify that this signal persisted even when factoring out potential covariate effects, we repeated a 2-stage procedure similar to classic TPOT, using resAdj TPOT with the adjustments described in Methods. Since resAdj TPOT transforms the problem from classification to regression, performance was measured by the coefficient of determination, as opposed to accuracy. Because of how resAdj TPOT operates, in stage 1 the Template used was *FeatureSetSelector-resAdjTransformer-Transformer-Regressor*; as the *resAdjTranformer* is required in order to make the adjustments. For the same reason, in stage 2, we had less flexibility and could not dispense of a Template (we used *resAdjTransformer-Transformer-Regressor*). In the first stage runs, out of the 197 FSs again the FS corresponding to PRC1 was the one most frequently selected, in 7 out of 43 successful runs. Albeit the best coefficient of determination in stage 2 was low (0.0023), the frequency with which the FS for *PRC1* occurred in stage 1 indicates presence of signal in this FS. We note that a permutation approach in the spirit of what we did for classic TPOT could not be applied here, because permuting the target column would have disrupted the relationship between target and covariates.

In order to better understand the drivers of the model in the best pipeline from stage 2, based on the FS consisting of the 28 SNPs mapping to the body, promoter, or enhancers of *PPP2R3A* and *PRC1*, we looked at permutation feature importance, both for classic and resAdj TPOT. The drivers for the best stage 2 pipeline from classic TPOT (illustrated in Fig. 3) were the SNPs rs4932178 and rs113028686. These two SNPs were also among the top 7 SNPs in the best stage2 pipeline from resAdj TPOT, together with rs116415933, rs139138366, rs8031684, rs11073964, rs35773450. The SNP rs4932178 resides in a putative enhancer for *PRC1* (in heart and fat), but is also within the promoter of *FURIN*, a gene expressed in vascular Endothelial Cells (ECs) and whose levels in ECs affect monocyte-endothelial adhesion and migration [26]. It has also been shown that *FURIN* inhibition reduces vascular remodeling and atherosclerotic lesion progression in mice [27]. Furthermore, this gene is among the prioritized causal CAD genes from [28] based on cumulative evidence from experimental and in silico studies. rs4932178 was also identified in GTEx (https://gtexportal.org/, v8) as an eQTL for *FES* in various tissues including coronary artery. Colocalization between CAD and expression association signals was observed for *FES* by [29]. rs113028686 is in the 5’-UTR of *PRC1* and is an eQTL of *FES* in various tissues including adipose, whole blood, and tibial artery (from GTEx). rs8031684, residing within an intron of *PRC1*, is an eQTL of *RCCD1* in adipose, aortic and tibial artery, as indicated in HaploReg v4.1 [30]. *RCCD1* is in the same subnetwork as *FURIN* for the CAD key driver *NGRN* identified in [31]. rs11073964, just upstream of *PRC1*, is also a missense mutation for *VPS33B* which is among 13 novel susceptibility loci for early-onset myocardial infarction identified in [32]. Among the remaining 3 SNPs (rs116415933, rs139138366, rs35773450), all intronic within the CAD druggable gene *PPP2R3A*, rs116415933 is reported in GTEx as eQTL for *IL20RB* in various tissues, including adipose and aortic and tibial artery. GeneCards [33] reports an association with the CAD phenotype for *IL20RB* (www.genecards.org).

**Fig. 3.**
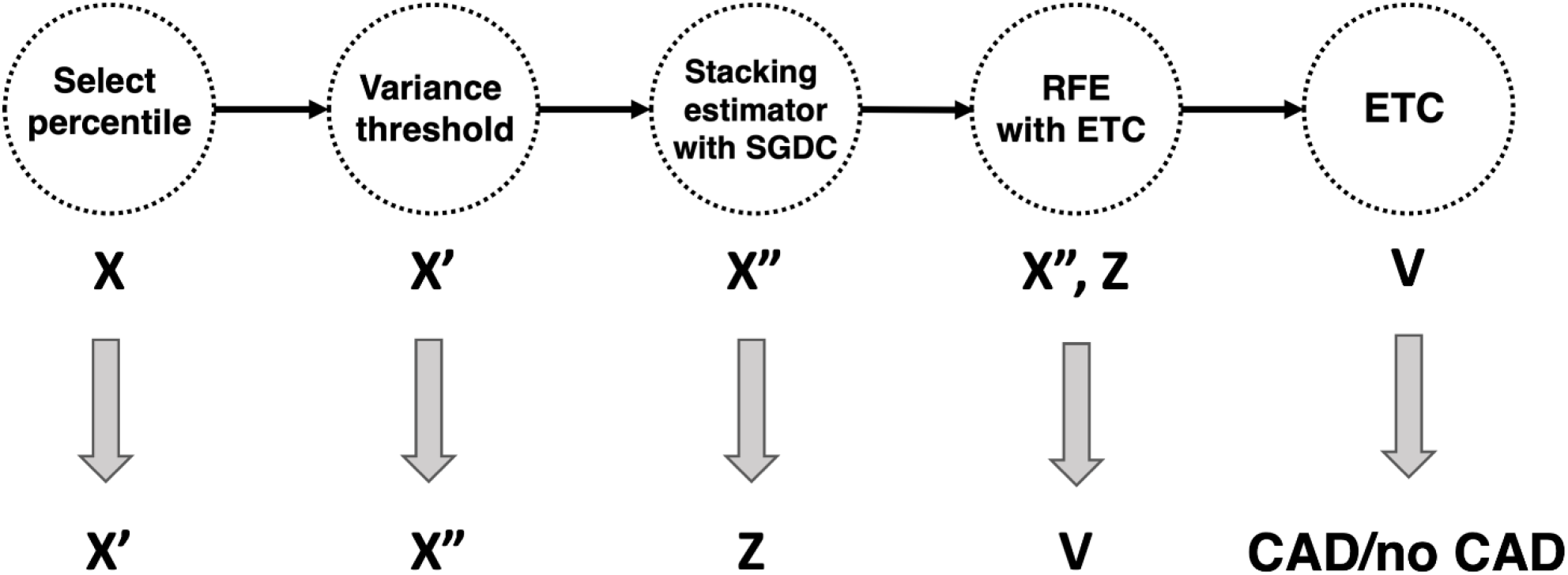
Outline of the best pipeline from the classic TPOT stage 2 runs. Select Percentile, Variance Threshold, and Recursive Feature Elimination (RFE) are feature selectors. Extra Trees Classifier (ETC) and Stochastic Gradient Descent Classifier (SGDC) are classifier estimators. The Stacking Estimator adds to its input features the results of applying the indicated estimator to those features.

Permutation importance measures the overall relevance of a feature to a model, i.e. how much the model relies on that feature, by examining how much shuffling the feature values increases the model error. However, especially when a model has limited predictive ability and heterogeneity exists among subjects, as it is typically the case with GWAS data, it is of interest to examine how each feature contributes to the individual predictions. With this in mind, we set to examine which features were driving the good predictions among the CAD cases using SHAP values, a game-theory based metric for explaining individual predictions [16]. We first examined the best pipeline from stage 2 of classic TPOT and computed the feature SHAP values for the 1489 testing CAD cases that were correctly classified. Fig. 4 shows the force plot for these subjects (force plots were introduced in [34]). Based on inspection of the force plot and Dunn indices for various *k* values, we used SHAP values to cluster these samples into four groups (sizes: 76, 156, 252, and 1005). We then ranked the features within each group by their average impact on model output across that group.

**Fig. 4.**
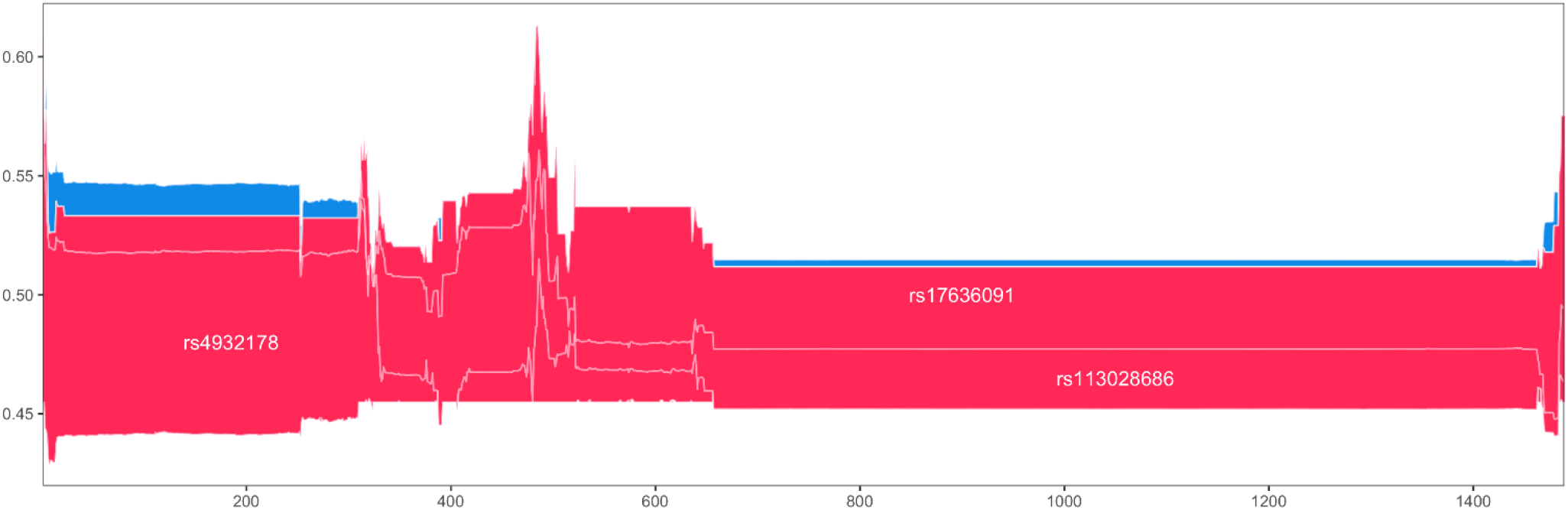
Multisample force plot for the 1489 correctly classified testing cases in the best stage 2 pipeline for classic TPOT. Explanations for these subjects are stacked horizontally, so the x-axis indicates the individuals. For each individual, the feature contributions to its prediction (probability of CAD) are shown along the y-direction, with features pushing the prediction higher in red, and features pushing the prediction lower in blue.

We observed that the same three features are driving the model output in all four clusters but with differing relevance (Fig. 5). These are two SNPs discussed above (rs4932178 and rs113028686) plus the rs17636091 SNP. The latter resides within an intron of *PRC1* and is reported by GTEx as eQTL, in various tissues, including adipose and artery (aorta and tibial), for both *RCCD1* and *VPS33B*, genes whose relevance to CAD has been discussed above. Clusters 1 and 3 are very similar with rs4932178 having a markedly higher average contribution to the model output, and rs17636091 contributing slightly more than rs113028686 on average in cluster 1 than in 3. For subjects in cluster 2, predictions are mostly driven by rs113028686, whereas for those in cluster 3, rs17636091 is the driver with rs113028686 not too distant of a second.

**Fig. 5.**
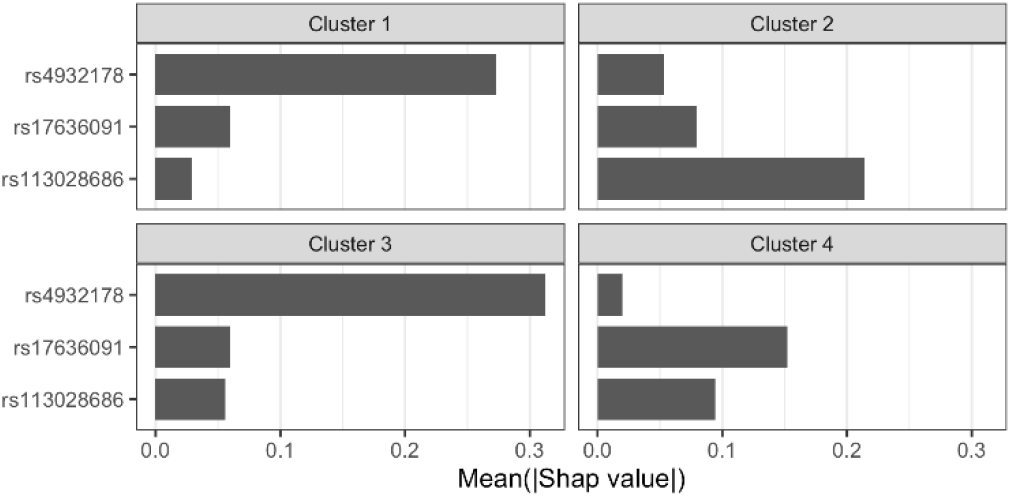
Feature rankings within the four SHAP value-based clusters for the correctly classified CAD testing cases in the best stage 2 pipeline from classic TPOT. The x-axis indicates the mean absolute SHAP value for the subjects in that cluster. Only the top 3 (out of 28) features are indicated as all remaining ones have negligible contributions.

We then proceeded with a similar approach for the best stage 2 pipeline from resAdj TPOT. We focused on the 250 CAD testing cases in the bottom quartile of the absolute difference between the covariate-adjusted observed and predicted outcomes. We note that these individuals are distinct from the 1489 correctly predicted using classic TPOT (overlap=19). Based on inspection of the force plot (Fig. 6) and Dunn indices for various *k* values, we clustered these subjects into six SHAP value-based groups (sizes 26, 5, 27, 13, 43, and 136).

**Fig. 6.**
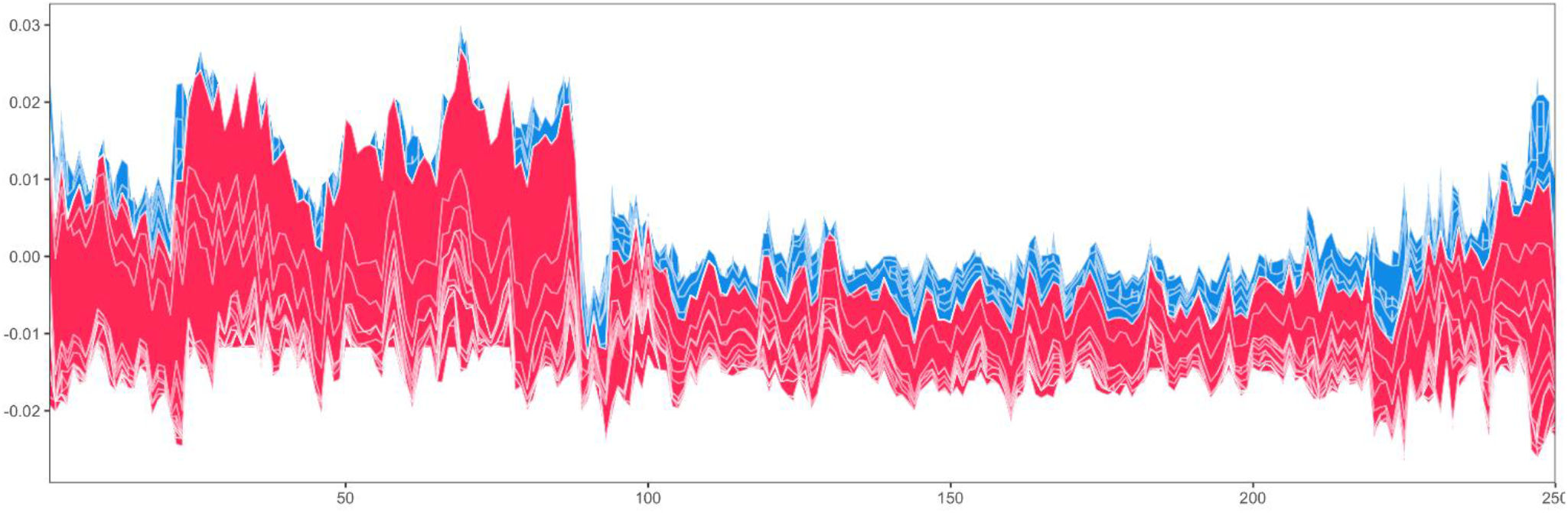
Multisample force plot for the 250 well classified testing cases in the best stage 2 pipeline for resAdj TPOT. Explanations for these subjects are stacked horizontally, so the x-axis indicates the individuals. For each individual, the feature contributions to its prediction are shown along the y-direction, with features pushing the prediction higher in red, and features pushing the prediction lower in blue.

In this case (Fig. 7), rs4932178 returns as the main driver to the model predictions for subjects in cluster 3. Moreover, this SNP is a strong driver in cluster 4 together with the top driver of that cluster, which is rs116415933, discussed above. The latter SNP is also the main driver in cluster 5, whereas in cluster 1 it is the main driver closely followed by rs188650245, an intronic SNP in *PPP2R3A*, indicated in GTEx as eQTL in several tissues, including cultured fibroblasts, for *IL20RB*, a gene discussed above. The latter SNP is also a top driver for cluster 6, but here with similar impact to rs139138366. Finally, cluster 2 is dominated by rs17636091, discussed above.

**Fig. 7.**
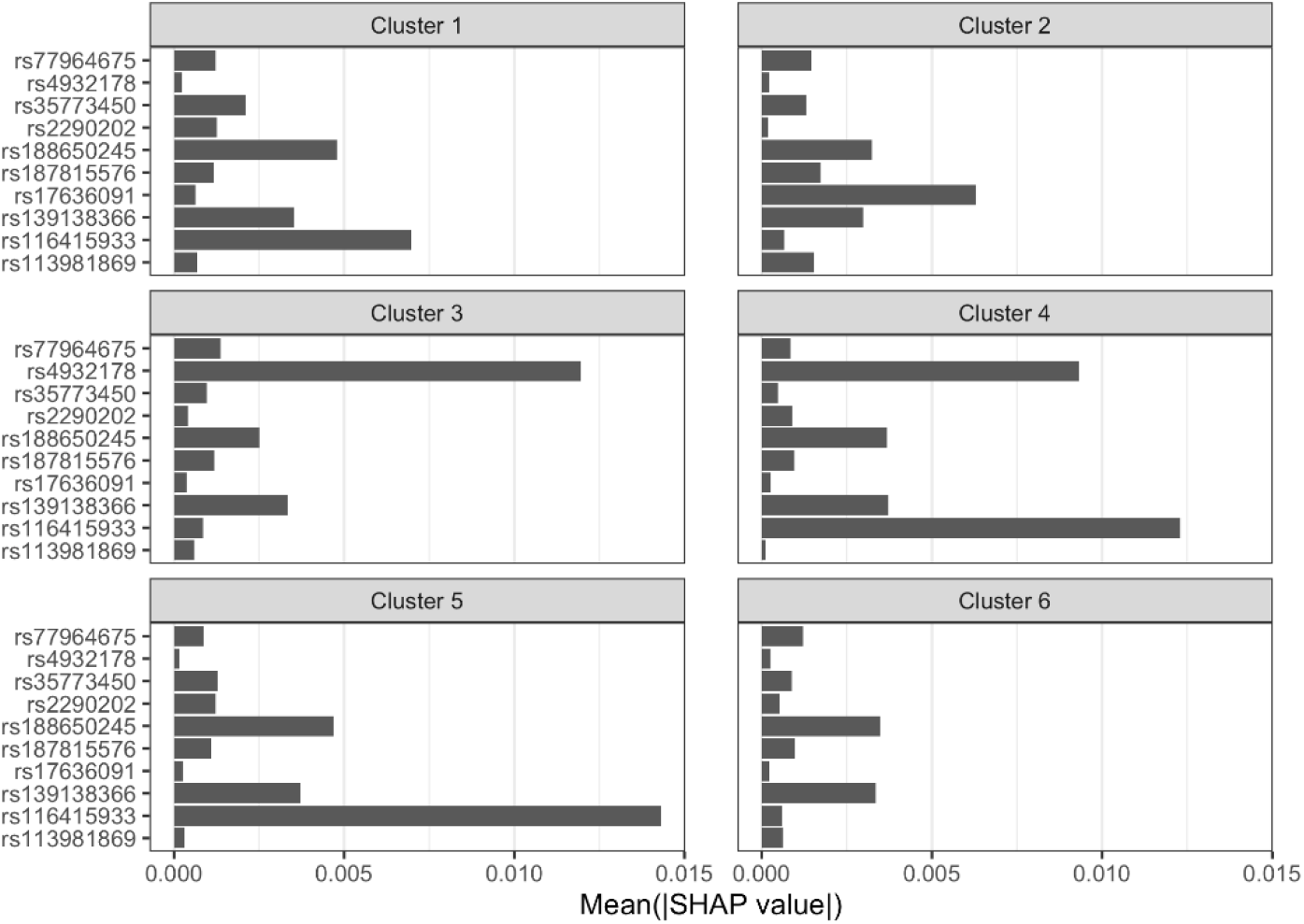
Feature rankings within the six SHAP value-based clusters for the well predicted CAD testing cases in the best stage 2 pipeline from resAdj TPOT. The x-axis indicates the mean absolute SHAP value for the subjects in that cluster. The features displayed are those in the union of the top 5 from each of the 6 clusters.

## 4 Conclusion and Discussion

In this work we employed a large-scale genotype data set for the CAD phenotype, derived from UKB, to assess the applicability of AutoML, specifically TPOT. Traditional GWAS analyses are based on univariate statistical approaches aimed at detecting main effects. ML approaches like TPOT enable investigation of SNPs as groups, embracing the possibility of both additive and epistatic effects. However, GWAS data sets present some unique challenges to AutoML as compared to other data types. First, the search space is very large, both in terms of number of observations (subjects) and features (SNPs). Moreover, if one is interested in comparing different feature sets, then the search space is indeed much larger than the search space for univariate analyses. All of this hinders computational feasibility. Second, the signal is weak and hard to detect, due to several inherent characteristics of this type of data, including small effect sizes of common variants and heterogeneity. Our baseline runs of TPOT, focused on features previously identified as the strongest CAD main effects, confirm this, with accuracies just above 55%.

In order to overcome the large search space obstacle, on the one hand, we randomly down-sampled the controls to equal the number of cases (about 20,000 each). We applied this down-sampling multiple times with different seeds before performing multiple TPOT runs. This down-sampling still retained a considerably large number of observations and at the same time eliminated the issue of having a highly unbalanced data set (even though the balanced accuracy score can be used in TPOT to deal with unbalanced data set). We also reduced the feature search space by employing biological filters where we integrated three resources: (1) results from a previous druggability prioritization study for CAD, (2) Hetionet integrated network, (3) tissue specific enhancer-promoter predictions derived from Roadmap Epigenomics data [35]. In addition, we employed various scorers to filter out potentially non-functional SNPs. Reducing the SNPs to be analyzed is a necessary but very delicate step. The idea is to utilize multiple lines of evidence to narrow down the feature space, whereby the goal is not to look for all possible signals of interest but to focus on a promising subset. There are many paths that can be taken to this end and in our case study we picked and followed one route. In general, for this type of filtering approaches, the risk is to discard too much and hence eliminate all potentially interesting features. The choice of suitable functional genomics data, public databases, and scoring algorithms bear a crucial weight into guiding the selection of SNPs and SNP groupings. As more functional genomics data become available as well as improved computational methods to extract from these more precise and relevant SNP-gene mappings, the application of TPOT and other AutoML to high-throughput genotype-phenotype data should become increasingly fruitful.

In spite of this reduction in search space size, the input data to our runs were still very large; thus, we had to set an upper bound of 100 for both the GP population and the GP number of generations. Moreover, for our permutation analyses, where the whole process had to be repeated for each permutation and we carried out 50 TPOT runs per permutation per stage, we had to limit the number of permutations to 20. Thus, improving TPOT runtime is an important area for further development so that runs on GWAS data can be on par with the typical settings used for other data types (e.g. 500-1000 generations and similarly sized populations in the GP). Improvements in run time would also enable increasing the number of permutations so to estimate p-values with small uncertainty.

A feature that turned out to be particularly useful for this scenario where accuracies were just above 50%, was the FSS which allowed us to specify that the pipelines being searched by TPOT in the first stage should all start with the selection of a feature set among a collection of sets of interest. With this, by examining the consistency with which a given FS was selected in multiple runs, we managed to identify a strikingly recurrent FS, comprising SNPs residing within the body, promoter, or enhancers of the ‘druggable’ *PPP2R3A* or its connected gene *PRC1*. To our knowledge, the latter gene has not previously been reported as being CAD relevant. Even though we cannot rule out the possibility that it represents a novel CAD gene, there are other possible explanations for the signal detected by TPOT. Indeed, when we examined permutation-based feature importance, we noted that several of the top SNPs relevant to models from the best stage 2 TPOT pipelines were not only in functional regions for *PRC1*, but also in functional regions for other genes (such as *FES, FURIN, RCCD1, VPS33B*, and *IL20RB*) with evidence for CAD relevance from previous studies. We note that permutation feature importance should not be overinterpreted in data sets like those derived from GWAS, where the predictive power is limited and heterogeneity is expected among the cases. In view of the latter, it is especially important to examine the feature contributions to model output on an individual basis. In this work, we computed SHAP values to cluster the testing cases with good predictions and examined the drivers of these predictions in different clusters. This approach underscores the case heterogeneity in our data set and provides an example of how to move towards precision medicine by utilizing metrics, such as SHAP values, which can help distinguish which features are relevant for which individuals. Our analysis of the SHAP value-based clusters also highlighted groups of subjects where the predictions were driven from SNPs associated to the genes with CAD relevance indicated above. Together, our findings corroborate that, despite the specific challenges presented by GWAS to TPOT, insights can be gained from applications of AutoML to this type of data, especially in combination with consistency measures (as provided by the FS recurrence analysis) and metrics aimed at facilitating model explanations such as SHAP.

## Acknowledgment

Funding for this study was provided by NIH grant LM010098. This research has been conducted using the UK Biobank Resource under Application Number 50978. We thank William La Cava for useful conversations about the SHAP python library. The corresponding author for this work is J.H. Moore.

## Supplemental material

**Supplemental Table 1**. SNPs in the *PPP2R3A*-*PRC1* feature set. For each SNP, its CADD, GWAVA and TraP scores are indicated together with the gene the SNP maps to and whether it resides within the extended gene body (B) or a putative enhancer (E).

